# Muscle stem cell activation in response to acute injury is promoted by transient exposure to neutrophil elastase

**DOI:** 10.1101/2025.02.26.640469

**Authors:** Christopher J. Clarke, Amy Stonadge, Ethan Aminov, Addolorata Pisconti

## Abstract

The molecular mechanisms underlying the activation of muscle stem cells in response to skeletal muscle injury are complex and only beginning to be elucidated since new tools were developed in the last decade. The immune response to tissue injury, often referred to as sterile inflammation, is a known key contributor to the mechanisms supporting muscle stem cell proliferation and differentiation in the muscle stem cell niche, but it is unclear whether muscle stem cell activation is also affected by the immune response. Here we show that neutrophil elastase, released in abundance during the first two days after muscle injury by infiltrating neutrophils, contributes to muscle stem cell re-entry into the cell cycle and commitment to the myogenic program. When neutrophil elastase is genetically ablated in mice, muscle stem cell activation is impaired, and muscle regeneration is delayed. Additionally, we identified cMet/mTOR and FGFR/ERK1/2 as two pathways activated by elastase in two different *in vitro* systems, via elastase-mediated cleavage and activation of pro-HGF and via elastase-mediated cleavage of extracellular matrix and release of bioactive FGF, respectively. While chronic exposure to neutrophil elastase impairs myoblast survival and differentiation, ultimately causing muscle regeneration defects, its transient presence in injured muscle during the first 24-48 hours post injury is beneficial and necessary. Thus, our results emphasize the importance of a tight regulation of the immune response for successful muscle regeneration, and provide strong evidence in support of a role for neutrophil elastase that is mainly to promote muscle healing in response to injury. Mechanistically, we highlight a key role for the cross-talk between neutrophils and muscle stem cells in muscle stem cell activation that involves neutrophil-derived elastase-mediated activation of mTOR and ERK1/2 via cMet and FGFR in muscle stem cells.

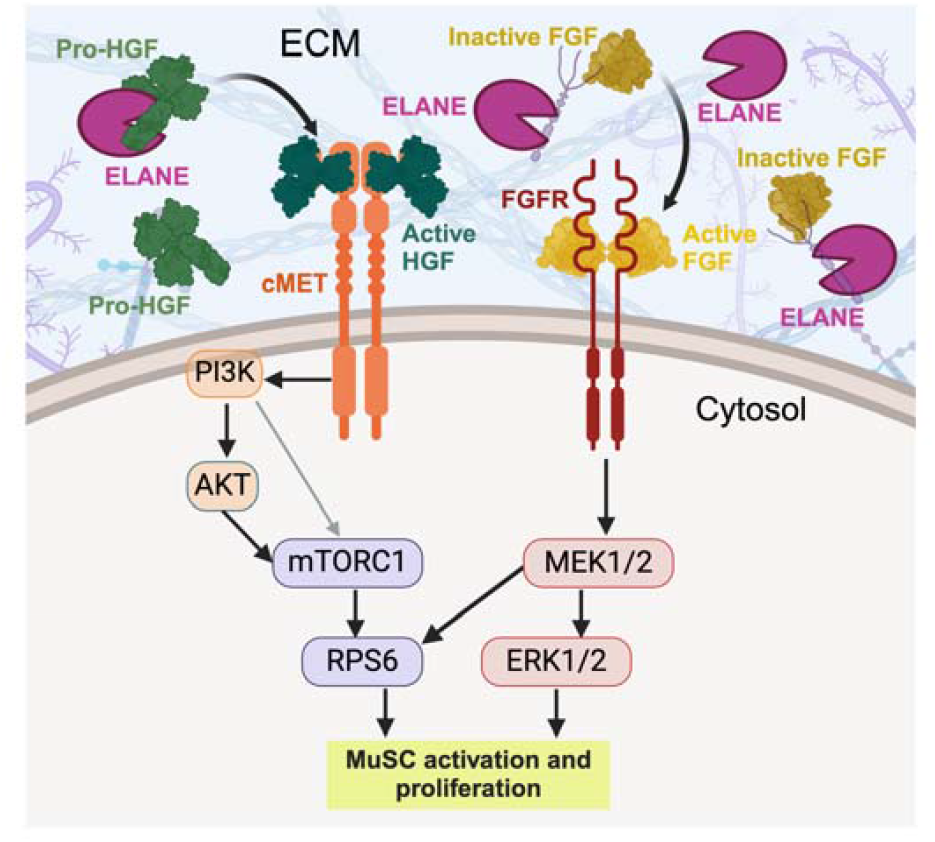

## INTRODUCTION

Regeneration of injured tissues is a complex biological process tightly regulated by cross-interactions between several cell types. Skeletal muscle regeneration in response to acute injury is achieved thanks to the extraordinary regenerative potential of a population of resident stem cells known as satellite cells, or muscle stem cells (MuSCs). In intact, healthy muscle, MuSCs are predominantly quiescent but poised to quickly become activated in response to injury, re-enter the cell cycle and extensively proliferate prior to differentiating and fusing to form new muscle fibers, or to repair existing damaged fibers^1-4^. MuSCs are supported in their function by a host of other cell types which together constitute the MuSC niche and include fibro-adipogenic progenitors (FAPs), endothelial cells, perivascular cells (mainly pericytes) and immune cells^2^. While MuSCs are indispensable for muscle regeneration, the stromal cells of the MuSC niche also play important roles as demonstrated by the fact that their ablation leads to impaired regeneration^5-10^.

The immune response to acute skeletal muscle injury is tightly regulated^10^. The first immune cells to infiltrate the muscle tissue, within a few hours after injury, are the neutrophils, which mount a robust acute phase response dominated by the release of several proteases and pro-inflammatory cytokines^11-14^. Neutrophil-derived proteases drive post-injury extracellular matrix remodeling, debris clearance. Neutrophils also coordinate the subsequent macrophage infiltration, which is characterized by a first wave of pro-inflammatory macrophages, that accompany and promote MuSC proliferation, followed by a second wave of anti-inflammatory macrophages, which promote myoblast differentiation^2,9,10^. When such exquisite temporal control of injury-induced inflammation is altered, for example in the case of muscular dystrophy-induced chronic inflammation, the immune response to tissue injury becomes dysregulated, causing more detrimental effects than beneficial ones^10,15-18^. Consistently, we have shown that chronic and dysregulated accumulation of neutrophils and especially the continuous secretion of neutrophil elastase (ELANE) is associated with impaired regeneration, chronic inflammation and fibrosis in dystrophic mice^19^. However, these observations raise the fundamental question: does neutrophil elastase play a beneficial role in muscle regeneration when its secretion is limited to a short interval after injury? Here we show that a short exposure to neutrophil elastase during injury-induced regeneration is indeed beneficial and it promotes muscle stem cell activation via cMet/mTOR and FGFR/ERK.

## RESULTS

### Transient exposure to ELANE promotes myogenesis in vivo and in vitro, in a MuSC autonomous manner

Neutrophils infiltrate injured muscles within a few hours after injury and are cleared within 2-3 days^19^. We have shown that in dystrophic muscle, the chronic accumulation of neutrophils is detrimental to muscle regeneration via the toxic effects of chronic exposure to neutrophil elastase^19^. However, during healthy regeneration the muscle tissue is only briefly exposed to neutrophil elastase and thus, we hypothesized that a short exposure to elastase activity is not detrimental to muscle regeneration but could instead be beneficial. To test this hypothesis, we administered elastase to murine myoblasts under the same culture conditions used in our previous work^19^ but this time for varying amounts of time. We observed that a longer, chronic treatment indeed impairs proliferation (Fig. 1A-B) as previously shown^19^, while a short, acute treatment promotes proliferation (Fig. 1A-B). Intriguingly, cells that were briefly exposed to elastase during the proliferation phase (Fig. 1A) also show enhanced differentiation later on, as measured by both fusion index (Fig. 1C-D) and myotube area (Fig. 1C, E).

**Figure 1.**
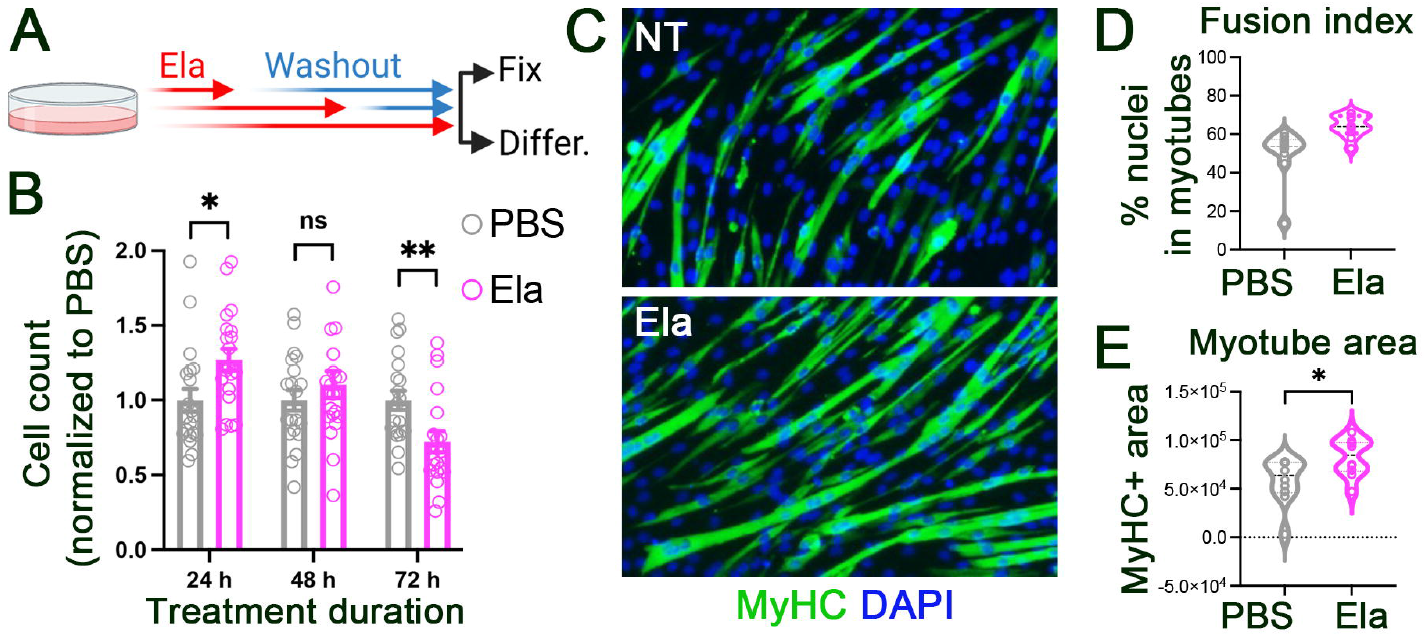
A brief exposure to elastase activity enhances myoblast proliferation and differentiation. **A)** C2C12 myoblasts were cultured as in Arecco et al., 2016^19^ and exposed to 0.5 U/mL of elastase for either 3 days (the same as in Arecco et al., 2016^19^) or shorter durations as indicated followed by wash out and incubation with vehicle control (PBS) for the remainder of the time. **B)** Three days after the beginning of treatments, cells were fixed and stained with DAPI to be scored for cell numbers. For each of three independent experiments the average of the +elastase condition (at least ten fields scored) was normalized to its internal vehicle (PBS) control (also at least 10 fields scored) and then the three averages combined and plotted (total N = at least 30) as individual point with the overall mean and S.E.M. indicated as bar and error bars, respectively. **C-E)** Cells were treated as in **A** but after three days in culture, instead of being fixed and stained for cell counts, they were induced to differentiate by serum withdrawal. Four days later, differentiated cultures were fixed and immunostained to detect skeletal muscle-specific myosin heavy chain (green) and counterstained with DAPI (blue). Panel C shows representative images of this experiment. **D)** The fusion index was calculated for at least 10 fields per experiment over three independent experiments and plotted as individual points for each individual score, while the distribution for each condition is depicted as a violin plot. **E)** The myotube size was calculated for at least 10 fields per experiment over three independent experiments and plotted as individual points for each individual score, while the distribution for each condition is depicted as a violin plot. *=p<0.05, **=p<0.01.

We next asked whether transient exposure to neutrophil elastase promotes myogenesis *in vivo* by measuring the muscle regenerative potential of neutrophil elastase knockout (*Elane*^*-/-*^) mice compared to that of wild type mice. Upon BaCl_2_ injury, the tibialis anterior muscles of wild type and *Elane*^*-/-*^ mice were harvested at 7 and 14 days after injury, or from uninjured mice (Fig. 2A). One week after injury, *Elane*^*-/-*^ muscles showed impaired regeneration, as measured by the percentage of centrally-nucleated fibers (Fig. 2B-C). One week later (two weeks after injury), regeneration was still delayed in *Elane*^*-/-*^ mice, which showed fewer and smaller regenerating fibers compared to wild type mice (Fig. 2D-E). The delay in regeneration was unlikely due to a defect in the immune response and “debris clearance”, since the macrophage burden was preserved in *Elane*^*-/-*^ mice at all time points analyzed (Fig. 2F), suggesting the timing of the immune response was unaltered. In contrast, loss of ELANE appears to directly impair MuSC commitment and differentiation in the early days after injury (Fig. 2G)

**Figure 2.**
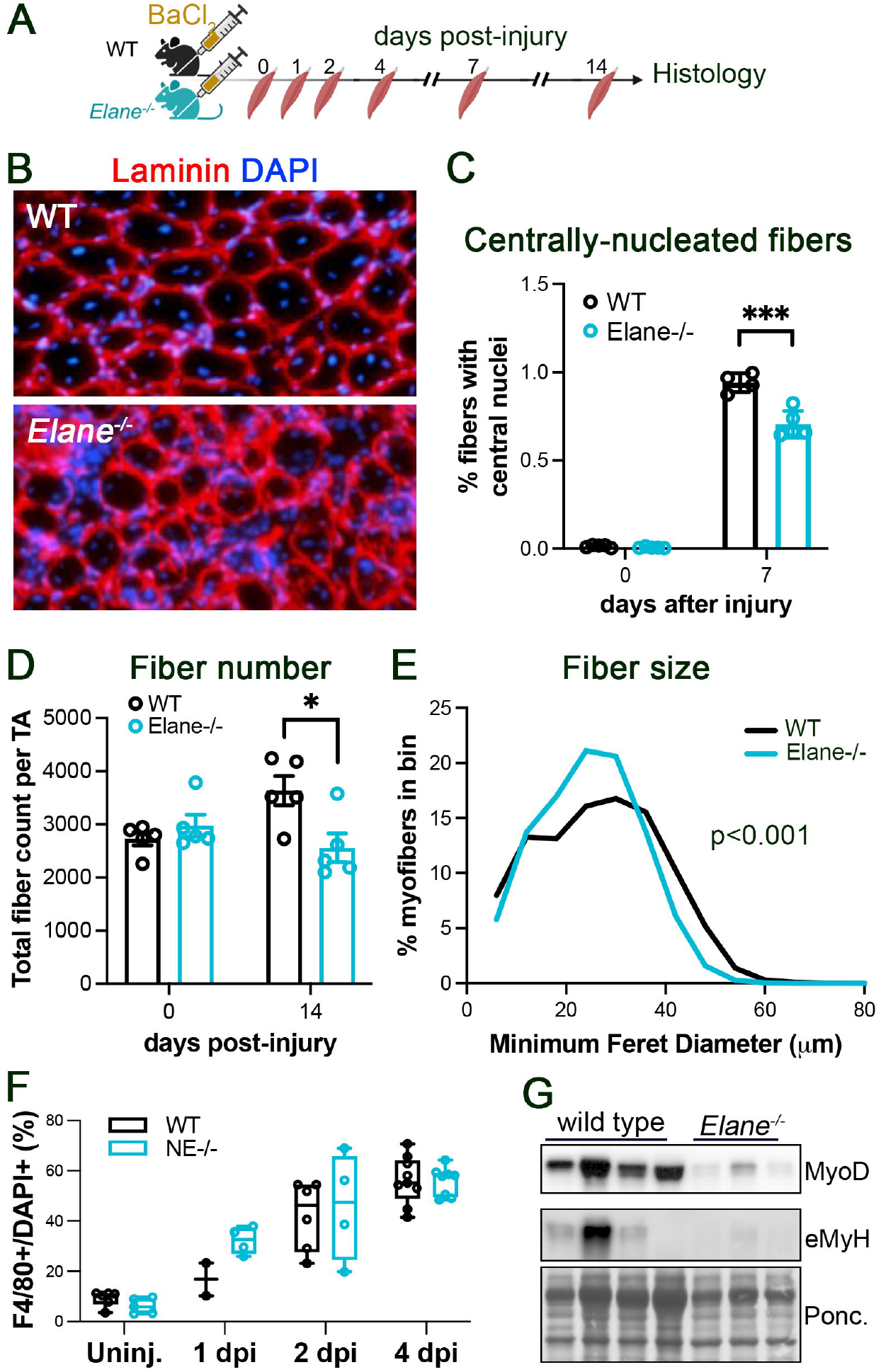
Neutrophil elastase (Elane) loss impairs MuSC activation *in vivo*. **A)** Wild type (WT) and *Elane*^*-/-*^ mice (at least 5 per time point and genotype cohort) received either a BaCl_2_ injection in the tibialis anterior (TA) muscle that caused extensive injury, or were left uninjured. Mice were then were euthanized at varying time points, and both their TA muscles dissected and frozen for further processing. **B)** Representative immunofluorescence of wild type and *Elane*^*-/-*^ TA muscle cross sections 7 days after injury, where laminin is depicted in red and DAPI in blue. **C)** Quantification of the percentage of centrally-nucleated myofibers in wild type (WT, gray) and *Elane*^*-/-*^ (teal) uninjured and injured mice, 7 days after injury (N = 5 per cohort). **D)** Total number of fibers in TA muscles of wild type (WT, gray) and *Elane*^*-/-*^ (teal) uninjured and injured mice, 14 days after injury (N = 5 per cohort). Distribution of myofiber size measured by the fiber minimum Feret diameter in wild type (WT, gray) and *Elane*^*-/-*^ (teal) uninjured and injured mice, 7 days after injury (N = 5 per cohort). **F)** Macrophage burden, measured as percentage of F4/80+ cells over total nuclei, in wild type (WT, gray) and *Elane*^*-/-*^ (teal) uninjured and injured mice, at various time points days after injury (N = 5 per cohort). **G)** Four days after injury wild type (WT, gray) and *Elane*^*-/-*^ (teal) injured TA muscles where homogenized and the MyoD and embryonic myosin heavy chain content measured as a proxy for the extent of regeneration. Ponceau staining of the membrane is shown as loading control. *=p<0.05, ***=p<0.001.

### Elastase activity promotes MuSC activation

The data shown so far suggest a role for ELANE in supporting myogenesis via acting directly on MuSCs and/or myoblasts. The timing of ELANE release during injury coincides with MuSC activation and the first cell division (Fig, 3A-B), suggesting that elastase activity might promote MuSC activation. Indeed, two days post injury, the percentage of activated MuSCs (i.e. that had re-entered the cell cycle) was decreased in *Elane*^*-/-*^ mice compared to wild type controls (Fig. 3C-D).

**Figure 3.**
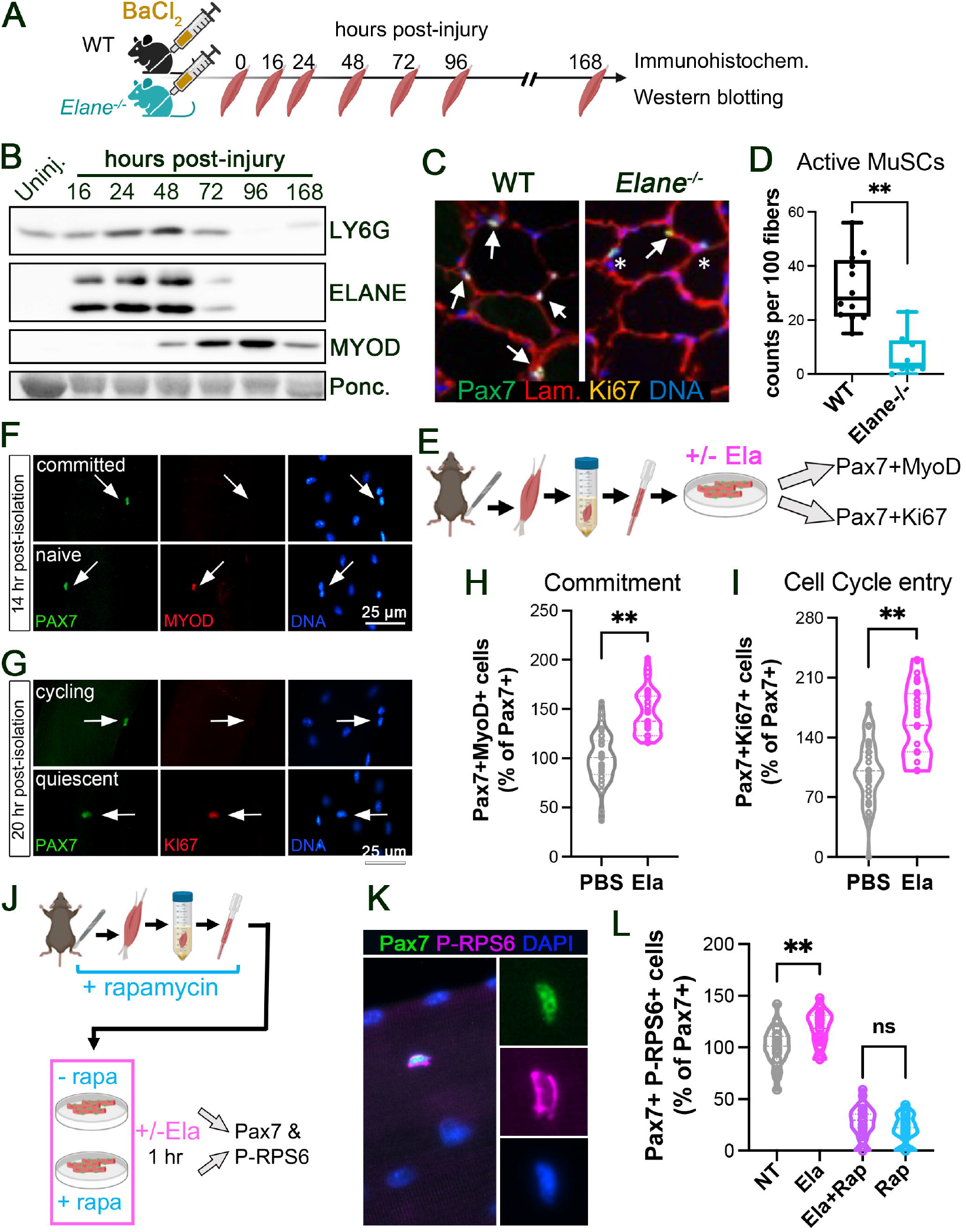
MuSC activation is impaired in *Elane*^*-/-*^ mice and enhanced by elastase treatment. **A)** Wild type (WT) and *Elane*^*-/-*^ mice (at least 5 per cohort) received either a BaCl_2_ injection in the tibialis anterior (TA) muscle that caused extensive injury, or were left uninjured. Mice were then euthanized at varying time points, and both their TA muscles dissected and frozen for further processing. **B)** Representative western blots of an injury time course where the neutrophil marker LY6G was detected alongside ELANE and MYOD, showing that neutrophils and ELANE are present in injured muscle during the first 48 hours, in concomitance with MuSC activation and beginning of commitment (note, after 48 hours MYOD increases by virtue of MuSC proliferation). **C)** Representative images of sublaminar cells expressing the MuSC marker Pax7 and the proliferation marker Ki67. Arrows are Pax7+Ki67+ MuSCs, asterisks are Pax7+Ki67-MuSCs. **D)** Quantification of C where 2 slides per animal, over a total of 4 animals, were scored and the average number of MuSCs per 100 myofibers plotted as mean +/-S.E.M. **E)** Diagram depicting the experimental design that yielded the data shown in F-I). **F-G)** Representative images showing examples of: F) committed (MYOD+) and non-committed (MYOD-) MuSCs (Pax7+), G) cycling (Ki67+) and non-cycling (Ki67-) MuSCs. **H)** Quantification of F where at least 10 fibers for each of three independent experiments were scored, each experiment was normalized to the vehicle treatment and then the individual datapoints of all three experiments were plotted as individual points. The violin plots depict the data distribution. **I)** Quantification of G where at least 10 fibers for each of three independent experiments were scored, each experiment was normalized to the vehicle treatment and then the individual datapoints of all three experiments were plotted as individual points. The violin plots depict the data distribution. **J)** Diagram depicting the experiment that yielded the data shown in K-L). **K)** Representative images showing an example of an early activated MuSC expressing phospho-RPS6. **L)** Quantification of (K) where at least 10 fibers for each of three independent experiments were scored, each experiment was normalized to the vehicle treatment and then the individual datapoints of all three experiments were plotted as individual points. The violin plots depict the data distribution.

To further test whether elastase activity promotes MuSC activation in a cell autonomous manner, we isolated muscle fibers with their associated MuSCs and treated them with either vehicle or elastase (Fig. 3E). After overnight incubation, the percentage of committed (MyoD+, Fig. 3F-H) and cycling (Ki67+, Fig. 3G-I) MuSCs in elastase-treated cultures was nearly double that of vehicle treated cultures.

One of the earliest pathways leading to MuSC activation in response to niche stimuli is the mTOR pathway^20^. Indeed, upon isolation, a large proportion of myofiber-associated MuSCs already show high levels of phospho-RPS6, which is directly downstream of mTOR kinase activity, unless the myofiber preparation is carried out in the presence of rapamycin at every step (Fig. 3J). Treatment with elastase significantly increases the percentage of phospho-RPS6+ myofibers in an mTOR-dependent manner (Fig. 3J-L), suggesting that elastase acts extracellularly on one of the signaling pathways that is transduced via mTOR.

### Elastase activity promotes MuSC activation via HGF/mTOR and FGF/ERK

The mTOR pathway is engaged by several niche borne signals, some of which have been described as early activators of MuSCs^20,21^. To dissect the molecular mechanisms underlying elastase-mediated activation of quiescent MuSCs we turned to a validated model of MuSC activation developed by Arora et al., where the murine myoblast cell line C2C12 is induced to undergo a quiescent (BrdU and Ki67 negative), uncommitted (MyoD negative) state by culturing the cells for two days in suspension (Fig. 4A and ^22^). Upon validation of this G0 synchronization model system (Supplementary Fig. S1), we induced cell cycle re-entry of G0-synchronized myoblasts (modeling MuSC activation) in the presence/absence of elastase overnight. Under these conditions, we observed both a time-dependent (Fig. 4B) and a concentration-dependent (Fig. 4C) positive effect of elastase on cell cycle re-entry. Crucially, the effect of elastase on promoting myoblast cell cycle re-entry was associated with AKT and RPS6 phosphorylation in a time-dependent manner (Fig. 4D), which further validated this model system as an appropriate one to dissect the molecular mechanisms underlying elastase-mediated activation of MuSCs via mTOR.

**Figure 4.**
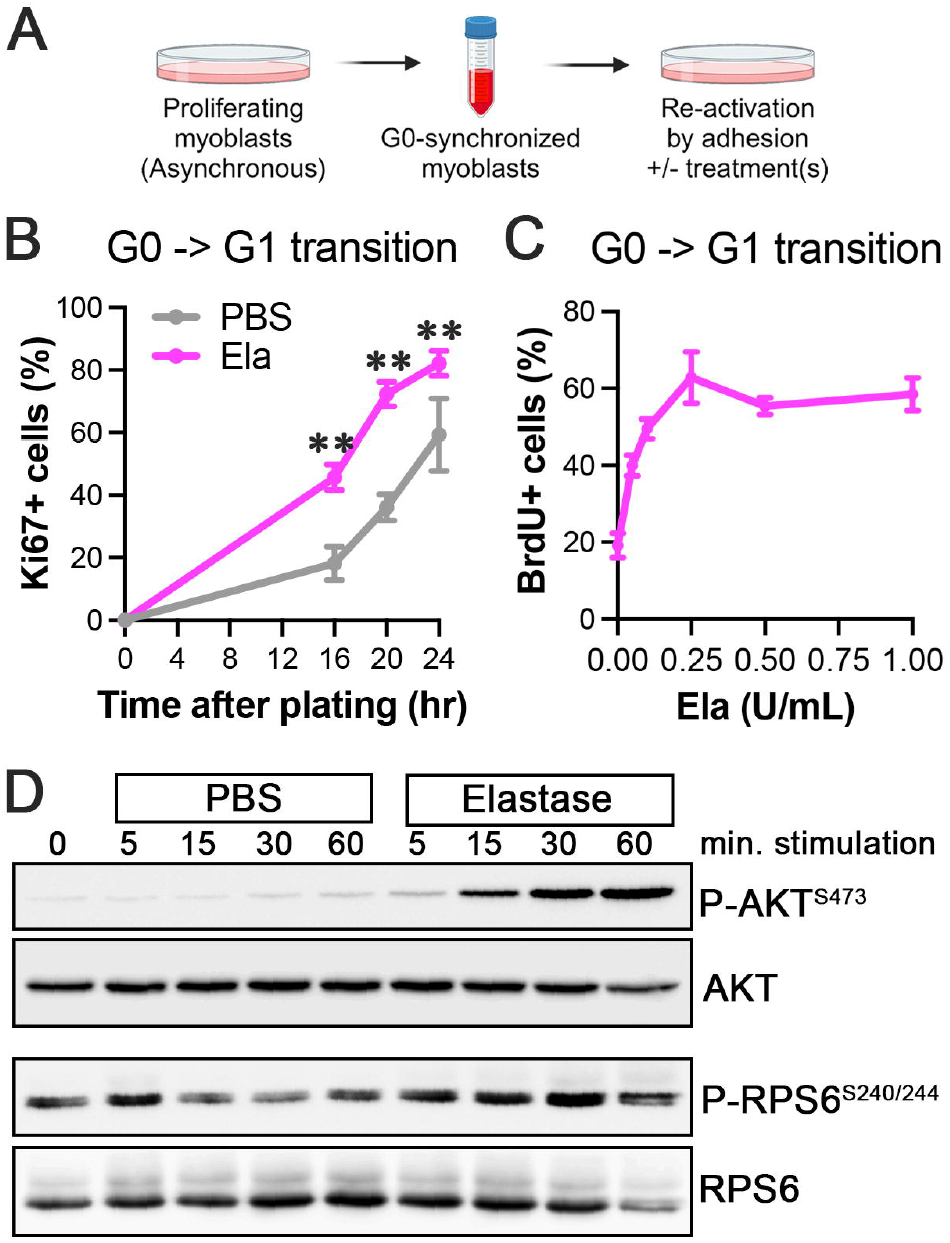
AKT/mTOR signaling is triggered by elastase in myoblasts transitioning from G0 to G1. **A)** Diagram depicting the experimental design that yielded the data shown in this figure. **B)** Time course of G0-synchronized myoblasts transition into the cell cycle (measured as % of cells expressing Ki67) in the presence (pink) or absence (gray) of elastase. **C)** Dose dependence of the effect of elastase on promoting G0-synchronized myoblast entry into S-phase (measured as % of cells incorporating BrdU. **D)** Western blotting of G0-synchronized myoblasts entering the cell cycle (3 hours after plating, thus at the time of G0-G1 transition) that were exposed to elastase for the indicated amount of time, detecting AKT phosphorylated in serine 473, total AKT, RPS6 phosphorylated on serine 240/244 and total RPS6. **=p<0.01.

As mentioned above, several signaling pathways that are transduced via mTOR have been shown to participate in MuSC activation and early proliferation, including HGF, EGF, FGF, IGF, PDGF and VEGF^21,23-32^. To gain insight into which external stimulus/i may be involved in elastase-mediated promotion of MuSC activation, we G0-synchronized myoblasts as previously described (Fig. 4A) and then, while releasing them back into the cell cycle, we treated them with either vehicle or elastase, alone or in combination with a range of growth factor receptor inhibitors (Fig. 5A). An elastase inhibitor was used as a positive control and the readout comprised AKT and RPS6 phosphorylation after 30 minutes of stimulation (Fig. 5B). Additionally, we studied phosphorylation of the same targets (RPS6 and AKT), also as a result of elastase treatment in the presence/absence of inhibitors specific for the kinases directly upstream of the targets of interest – i.e. PI3K upstream of AKT, AKT upstream of mTOR and mTOR upstream of RPS6 (Fig. 5C). With this approach we observed that both AKT and RPS6 are activated by elastase in a cMet (HGFR) and PI3K-dependent manner (Fig. 5B-C). Additionally, mTOR inhibition strongly prevented RPS6 phosphorylation (Fig. 5B-C). However, AKT inhibition did not prevent elastase-promoted RPS6 phosphorylation (Fig. 5B-C), raising the question: how are HGF/PI3K and mTOR/RPS6 linked if not via AKT? Intriguingly, MEK1/2 inhibition prevented elastase-promoted RPS6 phosphorylation but not AKT phosphorylation, suggesting a convergence of the MAPK pathway with the mTOR pathway downstream of AKT.

**Figure 5.**
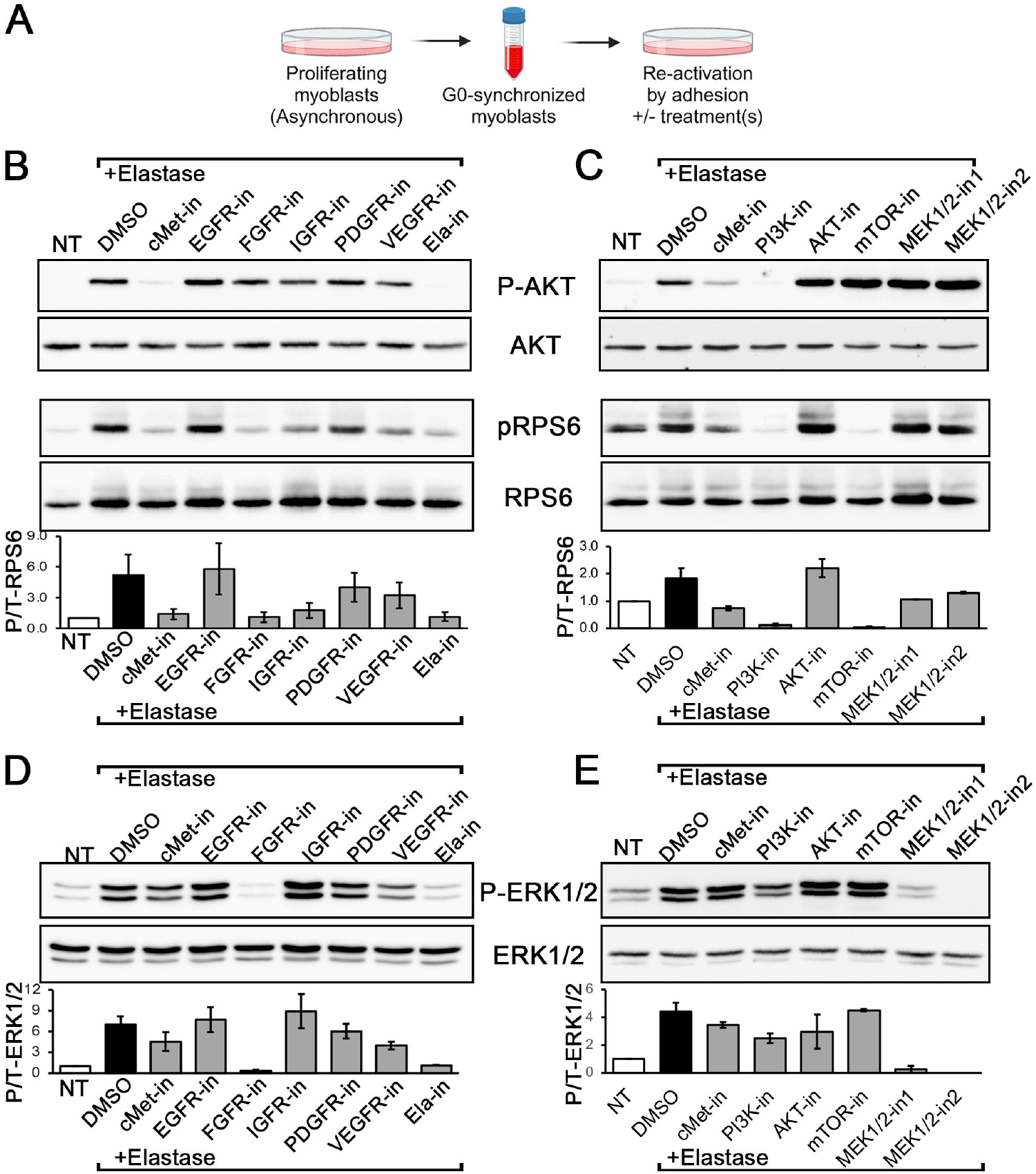
In response to elastase, myoblasts undergoing the G0-G1 transition trigger mTOR signaling via cMet, and ERK1/2 signaling via FGFR. **A)** Diagram depicting the experiment that yielded the data shown in this figure. **B-C)** Myoblasts were first G0-synchronized, then released back into the cell cycle via plating and 3 hours later exposed to either vehicle or elastase, the latter either alone or in combination with various inhibitors as indicated. Western blotting performed to detect AKT phosphorylated in serine 473, total AKT, RPS6 phosphorylated in serine 240/244 and total RPS6. Quantification of phospho-RPS6 normalized to total RPS6 is shown in the bar charts. **D-E)** Myoblasts that were first G0-synchronized, then released back into the cell cycle via plating, and 3 hours later exposed to either vehicle or elastase, the latter either alone or in combination with various inhibitors as indicated. Western blotting performed to detect ERK1/2 phosphorylated in threonine 202/204 and total ERK1/2. Quantification of phospho-ERK1/2 normalized to total ERK1/2 is shown in the bar charts.

All the growth factor receptors screened in the experiment described above can signal via both PI3K/AKT/mTOR and MEK/ERK (the MAPK pathway). Moreover, the mTOR and MAPK pathways downstream of growth factors are often coupled^33-35^. Thus, we speculated that elastase signaling can be transduced via both mTOR and ERK. ERK1/2 phosphorylation was strongly upregulated in response to elastase in a time-dependent manner (Supplementary Fig. S2A). When ERK1/2 phosphorylation was measured in response to elastase treatment and in the presence/absence of the various receptor inhibitors and kinase inhibitors described above, we observed a strong upregulation of ERK1/2 phosphorylation in response to elastase, that was FGFR and MEK1/2-dependent (Fig. 5D-E). Interestingly, elastase-dependent ERK1/2 phosphorylation was moderately inhibited by an AKT inhibitor (Fig. 5D-E), suggesting another level of coupling between the mTOR and MAPK pathway at the level of AKT-ERK1/2.

Lastly, we considered a potential role for elastase in regulating mTOR activity via interfering with adhesion signaling. Elastase enzymes cleave several ECM proteins and ECM receptors^19,36^ some of which, the integrins, can signal via mTOR. Moreover, growth factor and integrin signaling are often coupled^37^. Thus, we hypothesized that elastase may activate mTOR via activation of integrin signaling. However, we observed that elastase treatment decreases FAK phosphorylation in a time-dependent manner (Supplementary fig. S2B), suggesting that mTOR activation in response to elastase is FAK-independent.

### Elastase mediates pro-HGF activation and FGF liberation from the extracellular matrix

HGF is secreted as an inactive precursor (pro-HGF) which can be cleaved by several proteases to produce the bioactive HGF-alpha^38^. Crucially, the promiscuous cleavage site that yields HGF-alpha and beta also contains an elastase consensus site. To understand the underlying mechanism of elastase-mediated activation of cMet, we tested whether pro-HGF can be cleaved by elastase. Indeed, we verified that HGF-alpha release from pro-HGF is mediated by elastase in a dose-dependent manner (Fig. 6A) and is inhibited by an elastase inhibitor (Fig. 6A).

**Figure 6.**
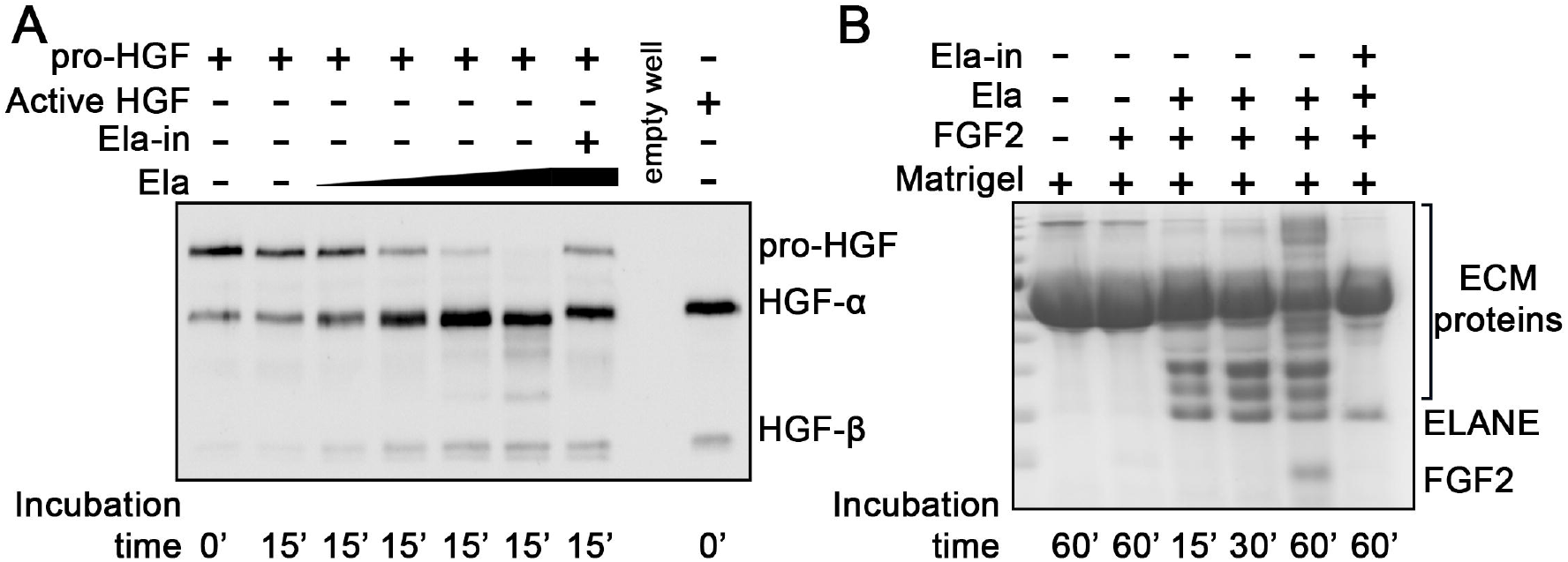
Elastase activity mediates HGF and FGF proteolytic release from pro-HGF and the ECM, respectively. **A)** Pro-HGF was incubated with either vehicle (PBS) or increasing concentrations of elastase for 15 minutes, in the absence/presence of an elastase inhibitor. The relative amounts of total pro-HGF, HGF-alpha and HGF-beta were assessed by gel electrophoresis through Coomassie staining and using a fully bioactive HGF (fully cleaved into alpha and beta subunits) as control. **B)** FGF was encapsulated in Matrigel and exposed to elastase for varying amount of time in the absence/presence of an elastase inhibitor. The relative amounts of free FGF and ECM proteins were assessed by gel electrophoresis through Coomassie staining. As the incubation time progresses, the ECM becomes more degraded and a band corresponding to FGF appears at 60 minutes incubation.

FGFs are secreted as bioactive molecules that do not require proteolytic activation, however their activity is inhibited by their binding to extracellular matrix (ECM) components, especially proteoglycans^39-41^. To test whether elastase activates FGF signaling by cleaving ECM proteins and therefore promoting FGF release from the matrix, we encapsulated FGF2 in Matrigel and incubated the Matrigel plugs with either vehicle or elastase, with or without an elastase inhibitor, for varying amounts of time. As the incubation time increased, we observed a progressive increase in ECM degradation products and a concomitant increase in free FGF2, which was fully inhibited by the elastase inhibitor (Fig. 6B).

### cMet/mTOR and FGFR/ERK1/2 are both necessary but not sufficient to mediate elastase-induced MuSC activation

Once the molecular mechanisms mediating elastase activity during the G0 to G1 transition had been dissected, we sought to validate their physiological relevance to MuSC activation. First, we used the same G0 to G1 transition model that had enabled us to identify HGF/cMet/mTOR, FGFR/ERK1/2 and FGFR/mTOR as transducers of elastase activity. We again synchronized myoblasts in G0 and induced them to de-commit (downregulate MyoD), by culturing them for two days in suspension, after which we released them into the cell cycle in the presence/absence of elastase and/or various relevant pathway inhibitors (Fig. 7A). Interestingly, both the cMet inhibitor and the FGFR inhibitor completely abolished elastase-induced G0 to G1 transition (Fig. 7B). Consistently, both an mTOR inhibitor and a MEK inhibitor completely abolished elastase-induced G0 to G1 transition (Fig. 7C).

**Figure 7.**
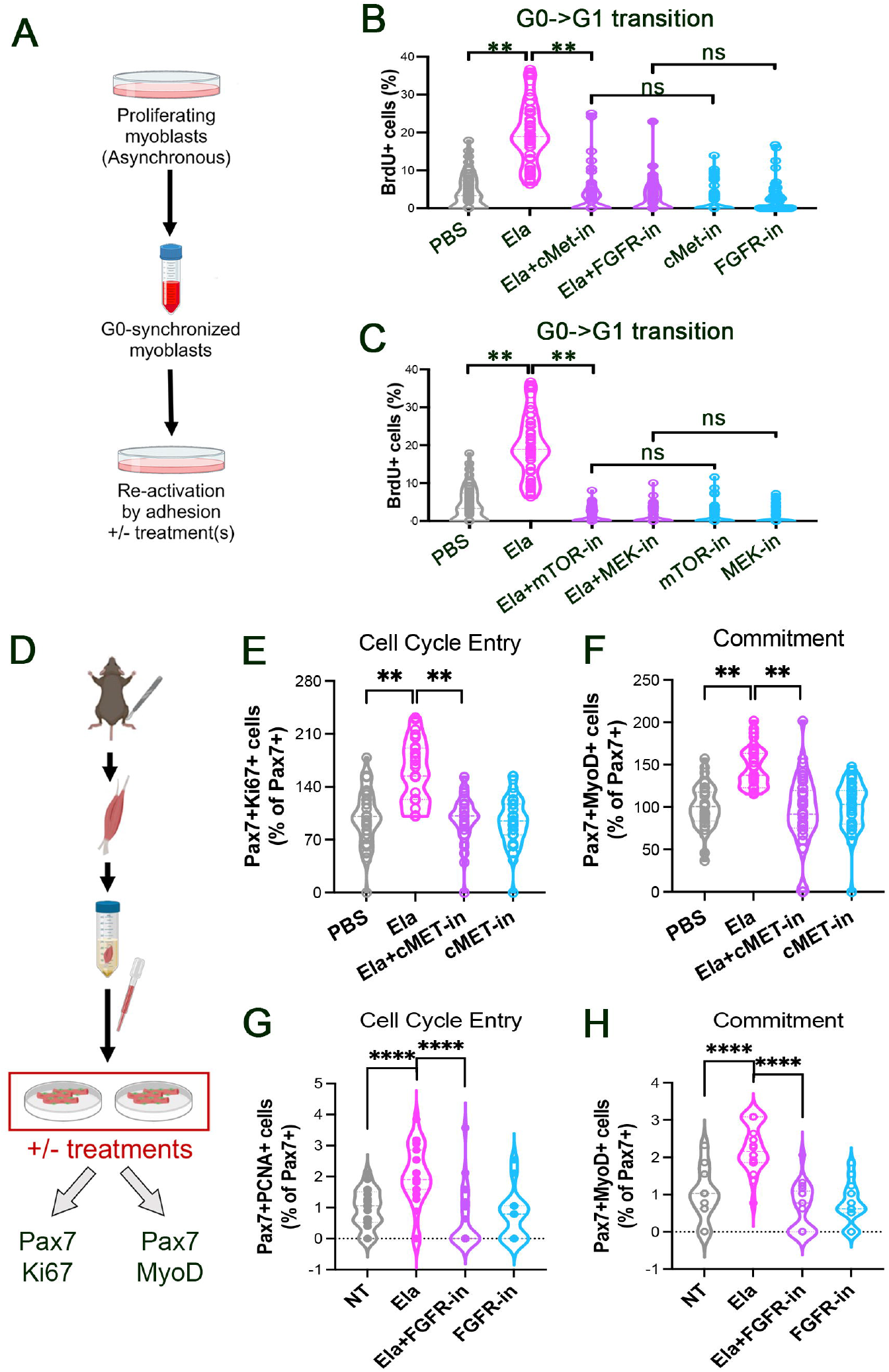
MuSCs activation is promoted by elastase via cMet/mTOR and via FGFR/ERK1/2. **A)** Diagram depicting the experimental design that yielded the data shown in B-C). **B-C)** G0-G1 transition in myoblasts exposed to vehicle or elastase, the latter alone or in combination with cMET or FGFR (B), mTOR or MEK1/2 (C) inhibitors, as indicated. At least 10 fields for each of three independent experiments were scored, each experiment was normalized to the vehicle treatment and then the individual datapoints of all three experiments were plotted as individual points. The violin plots depict the data distribution. **D)** Diagram depicting the experiment that yielded the data shown in E-H). **E-F)** G0-G1 transition (E) and myogenic commitment (F) in myofiber-associated MuSCs exposed to vehicle or elastase, the latter alone or in combination with a cMET inhibitor. **G-H)** G0-G1 transition (G) and myogenic commitment (H) in myofiber-associated, primary MuSCs exposed to vehicle or elastase, the latter alone or in combination with elastase or elastase and an FGFR inhibitor.

To further assess the role of these pathways in elastase-mediated MuSC activation, we next used a more physiological model where the G0 to G1 transition and myogenic commitment of primary MuSCs were assessed in the context of their native, myofiber niche (Fig. 7D). Again, we observed that all tested inhibitors were capable of fully abolishing elastase-induced MuSC activation (Fig. 7E-H).

## DISCUSSION

We have shown here that elastase activity promotes MuSC activation *in vivo* and *in vitro*, leading to an overall promotion of myogenesis. This effect of elastase in myogenesis is mediated by at least HGF/cMet through mTOR, and by FGFR through both ERK1/2 and mTOR.

From the physiological and functional point of view, our findings establish neutrophil infiltration, and especially secretion of ELANE, as an important player in MuSC activation in response to injury. In sterile inflammation, neutrophils are traditionally thought to exacerbate the extent of injury and further degrade the site of injury in preparation for clearance of the tissue debris my macrophages^42^. However, it is not surprising that the system has evolved such that tissue degradation via secreted proteases also participates in stem cell activation, eventually promoting tissue regeneration. Indeed, the idea that neutrophils also participate in tissue healing has been recently investigated in various settings^43-46^. Here we provide strong evidence in support of a role for neutrophil elastase that is mainly to promote muscle healing in response to injury, by showing that mice where ELANE has been genetically ablated show delayed regeneration. Moreover, we dissect the molecular mechanisms underlying the effects of elastase activity on MuSCs.

We chose to investigate the cMet and FGFR pathways as inhibition of either of these receptors yielded the most dramatic effects on two central effectors in MuSC activation: AKT and ERK1/2. Our results clearly indicate engagement of both mTORC1 and ERK1/2 via cMet and FGFR downstream of elastase activity. However, AKT, albeit engaged by elastase-mediated activation of cMet, does not appear to be involved in mTORC1 activation by either cMet or FGFR in response to elastase, since its inhibition does not affect RPS6 phosphorylation. In contrast, MEK1/2 inhibition does lead to a reduction in RPS6 phosphorylation, a direct measure of mTORC1 activity. There are two possible explanations for this observation. One is that mTORC1 is exclusively activated by elastase in an AKT-independent, ERK1/2-dependent manner. The other is that AKT does activate mTORC1, but the effect of AKT inhibition on mTORC1 activity is masked by a compensating direct phosphorylation of mTOR and/or p70S6 (also upstream of RPS6 phosphorylation) by ERK1/2, both of which have been described^47,48^. Another piece of evidence for AKT and MEK1/2 pathways being redundant at activating mTORC1 is that inhibition of PI3K, which is upstream of both, completely prevented elastase-induced RPS6 phosphorylation. However, this observation could also indicate that another PI3K-regulated effector pathway is responsible for mTOR/RPS6 activation. The identity of this alternate pathway is unclear but could be a pathway involving PDK1, a PI3K-activated master kinase that directly phosphorylates and activates RPS6^49^. Lastly, it is worth noting that the increase in AKT phosphorylation caused by MEK1/2 and mTORC1 inhibition, is likely due to loss of mTORC2 inhibition by mTORC1, as mTORC2 phosphorylates AKT^50^. Indeed, we observe this phenomenon also with direct AKT inhibition, which causes increased mTORC2-mediated AKT phosphorylation as a consequence of loss of AKT/mTORC1 activity.

One fascinating aspect of our results is that MEK and mTOR are both necessary, and neither is sufficient, for elastase-mediated MuSC activation. ERK signaling is mainly related to transcription and commitment to DNA synthesis in G1^51^, while mTOR signaling mainly supports cell growth, protein translation and lipid biosynthesis. Thus, it is plausible that the system has evolved such that both complementary pathways must be recruited to achieve full MuSC activation. Our cell assay data demonstrate a dependence on mTOR and ERK1/2 signaling to efficiently allow G0 to G1 progression, as inhibition of either pathway prevents G1 progression of G0-synchronized myoblasts. This implies that ERK1/2 activation, largely mediated by FGFR activation, promotes cell cycle re-entry through an mTOR-independent pathway. However, this pathway must be activated in combination with mTOR, as inhibition of either, in isolation, prevents cell cycle re-entry. Thus, while here we have attempted to shed light on the mechanisms underlying elastase-mediated promotion of MuSC activation, it is clear that none of these signaling events happen in isolation.

In conclusion, we have shown, using *in vivo, ex vivo* and *in vitro* approaches, that ELANE is secreted by infiltrating neutrophils in injured muscle during the early hours after injury, where it supports MuSC activation by cleaving and activating HGF, as well as by promoting release of FGF from the extracellular matrix. Taken together our data offer compelling evidence to the importance of a tightly regulated immune response to muscle injury. Moreover, our work is the first to describe in mechanistic detail the crosstalk between neutrophils and MuSCs during muscle regeneration, and identify ELANE as an important player.

## MATERIALS AND METHODS

### Cell Culture

#### C2C12 myoblasts

were cultured as previously described^19^ in DMEM (high glucose with pyruvate and GlutaMax) + 10% FBS (Invitrogen) + 1% Penicillin/streptomycin (growth medium). Cells were maintained at 40-70% confluence, thus passaged every other day.

#### Myoblast differentiation

Myoblast differentiation was induced by decreasing the serum to 2% when cells reached 80% confluence. The differentiation index was measured as the percentage of all total cells that expressed skeletal muscle-specific myosin heavy chain. Myotube area was measured with a bespoke written macro in ImageJ/Fiji as described before^52^.

#### Myofiber cultures

were obtained and maintained as previously described^53^. Briefly, the gastrocnemius muscles of wild type mice were digested for 1.5 hours at 37 °C, in 400 U/mL of collagenase type I dissolved in F12 medium supplemented with 2 mM CaCl_2_ (F12C), gently inverting every 15 minutes. At the end of the digestion period 3 volumes of F12C + 15% horse serum were added dilute the collagenase and individual myofibers were picked with a 3 mm glass Pasteur pipette after fire-polishing the tip. Fibers were moved into fresh F12 + 15% horse serum and washed two more times by serially moving them to fresh (warmed to 37 °C) F12C + 15% horse serum. Cultures were maintained in suspension bathed in F12 + 15% horse serum with or without treatments.

#### Drug treatment

For most of the experiments, elastase purified from porcine pancreas (Sigma) was used upon validation of its effects of myoblasts and myofiber-associate muscle stem cells, which were found to be identical to those of elastase extracted from human leukocytes (Sigma). In both cases, elastase was resuspended in cell culture grade phosphate buffered saline (PBS) and therefore PBS was used as vehicle in control experiments. Receptor and kinase inhibitors were dissolved in dimethysulfoxide (DMSO) and therefore DMSO was used as their vehicle in corresponding negative control experiments. The inhibitors used are listed in table 1.

**Table 1:** List of inhibitors used, their target, source and concentration.

| Inhibitor | Target | Source | Concentration used |
| --- | --- | --- | --- |
| AZD5363 | Akt | Tocris Bioscience | 1 uM |
| Crizotinib | cMet | Selleckchem | 1 uM |
| Erlotinib | EGFR | Selleckchem | 1 uM |
| Elastatinal | Elastase | Sigma | 100 ug/mL |
| Infigratinib | FGFR | Prof. David Fernig | 1 uM |
| AG1024 | IGFR | Sigma | 10 uM |
| Trametinib | MEK1/2 | Selleckchem | 1 uM |
| Rapamycin | mTOR | Selleckchem | 1 uM |
| Imatinib | PDGFR | Selleckchem | 1 uM |
| Wortmanin | PI3K | Sigma | 0.1 uM |
| Pazopanib | VEGFR | Selleckchem | 1 uM |

### Mice

*Elane*^*-/-*^ mice (generated on the C57Bl/6 background), were obtained from Jackson Laboratories, housed in a pathogen-free facility at the University of Liverpool, UK and used in accordance with the Animals (Scientific Procedures) Act 1986 and the EU Directive 2010/63/EU and after local ethical review and approval by Liverpool University’s Animal Welfare and Ethical Review Body (AWERB). Age-, sex- and background-matched wild type (C57Bl/6) controls were purchased from Charles River UK and housed for at least 2 weeks in the same facility as *Elane*^*-/-*^ mice before use.

#### Injury model

The right tibialis anterior (TA) muscle of male *Elane*^*-/-*^ and wild type mice aged 12-15 weeks was injured by direct intramuscular injection of 50 μL of 100mM? BaCl_2_ using an insulin syringe fitted with a G31 needle. The left contralateral TA muscle was left uninjured. Both TAs were dissected at various time points after injury upon humanely euthanizing the mice and were either snap-frozen in liquid nitrogen for subsequent protein extraction or embedded in Optimal Cutting Temperature (OCT) compound and frozen over a block of isopentane previously cooled to freezing temperature in a liquid nitrogen bath.

### Immunofluorescence

#### Immunofluorescence of Cells

Cells were fixed with 4% paraformaldehyde (PFA, prepared in PBS, pH 7.4), which was washed off three times with PBS prior to being permeabilized with 0.2% Triton X100 (prepared in PBS) for 10 minutes and blocked with 10% horse serum for 1 hour. Antibody incubation was usually carried out overnight at 4 °C in a humidified, dark chamber. Antibodies were diluted in PBS + 1% horse serum and were: mouse anti-myosin heavy chain (MF20 clone, DSHB), 1:100; rabbit anti-laminin (Sigma), 1:1000; rat anti-laminin alpha-2 (Sigma), 1:500; rabbit anti-Ki67 (abcam), 1:400; mouse anti-Pax7 (DSHB, purified) 1:100; mouse anti-MyoD (BD Biosciences, clone 5.8A), 1:100; rat anti-BrdU (Rockland), 1:100; rabbit anti-phospho-RPS6 (Cell Signaling Technologies), 1:100; rabbit anti-F4/80 (Cell Signaling Technologies), 1:100. Following washes with PBS, cells were incubated for 1 hour at room temperature with secondary antibodies, diluted at 1:500 in PBS + 5% horse serum, which were all from Invitrogen/Molecular Probes and conjugated to AlexaFluor-488 or AlexaFluor-555 or AlexaFluor-647, as needed. Lastly, cells were counter-stained with DAPI (Invitrogen) at a final concentration of 1 μM, diluted in PBS. Cells were usually imaged in their PBS bath on an EVOS M5000 microscope.

#### Immunofluorescence of muscle sections

Cryosections of injured and contralateral uninjured, or never injured TA muscles were obtained on a cryostat (Thermo Fisher) set a 10 μm thickness, -20 °C. Sections were permeabilized with PBS + 0.2% Triton X100 and the rest of the immunofluorescence carried out the same way as already described above for cells, with the only difference that at the end, slides were mounted with anti-fade VectaShield mounting medium (Vector Labs) and a thin glass coverslip (Fisher Scientific). Images were acquired on an EVOS M5000 microscope.

### Muscle morphometry

Percentage of centrally-nucleated myofibers, and myofiber size and numbers were measured using the functions “CNF” and “Properties”, respectively, of the software SMASH^54^. At least ten images per animal were analyzed for at least 5 animals per cohort.

### G0 synchronization of myoblasts

C2C12 myoblasts were synchronized in G0 as previously described^22^. Briefly, proliferating cells were trypsinized, resuspended in Methocel™ supplemented growth medium and incubated for 48 hours in a cell incubator (at 37 °C, in a 5% CO_2_ atmosphere). At this point, Methocel™ was diluted with PBS and the cells gently pelleted by centrifugation.Pelleted cells were then washed with PBS and resuspended in growth medium (DMEM + 10% FBS + 1% penicillin/streptomycin) and seeded on uncoated tissue culture dishes in the presence or absence of various treatments.

### Western blotting

Quadriceps from wild type and *Elane*^*-/-*^ mice were homogenized in RIPA buffer (150 mM NaCl, 50 mM Tris-HCl, pH 7.5, 1.0% IGEPAL, 0.1% SDS, 0.5% sodium deoxycholate) supplemented with a protease inhibitor cocktail (Complete, *Roche*) and phosphatase inhibitors (1 mM Na_3_VO_4_ + 1 mM NaF) using a VelociRuptor homogenizer (Scientific Laboratory Supplies), followed by incubation on ice for 30 minutes, and then cleared by centrifugation at 17,000 xg for 10 min at 4 °C. Western blotting was performed as previously described^55^. The antibodies used were: mouse anti-MyoD1 (BD Biosciences, clone 5.8A) at 1:500, mouse anti-eMyHC (F1.652 clone, DSHB) at 1:1000, rabbit anti-phospho-AKT(Ser473) (Cell Signaling Technology) at 1:1000, rabbit anti-phospho-RPS6(Ser240/244) (Cell Signaling Technology) at 1:1000, rat anti-phospho-ERK1/2(Thr202/204) (Cell Signaling Technology) at 1:1000, rabbit anti-AKT (Cell Signaling Technology) at 1:2000, rabbit anti-RPS6 (Cell Signaling Technology) at 1:2000, rabbit anti-ERK1/2 (Cell Signaling Technology) at 1:2000. Secondary anti-mouse and anti-rabbit antibodies, conjugated to horseradish peroxidase, were from Cell Signaling Technology and all used at 1:5000. Enhanced chemiluminescence (ECL) was produced on the probed membranes using an ECL kit from Amersham.

#### Densitometry analysis

ECL images were acquired on an Amersham ImageQuant™ Western Blot imaging system at non-saturating exposure times and band intensity quantified using the *Gel Analysis* function in ImageJ/Fiji.

### Statistical analysis

For all experiments, unless otherwise specified (e.g. for RNA-sequencing) the software Graphpad was used to carry out statistical analysis. In all cases, single pairwise comparisons were analyzed with an appropriate statistical test upon testing for normality: datasets that passed the normality test were analyzed using a parametric test (usually t-test with Welch’s correction for non-equal variance). Datasets that did not pass the normality test were analyzed with a non-parametric test, usually Mann-Whitney.

