## Supplementary figures for "Muscle stem cell activation in response to acute injury is promoted by transient exposure to neutrophil elastase"

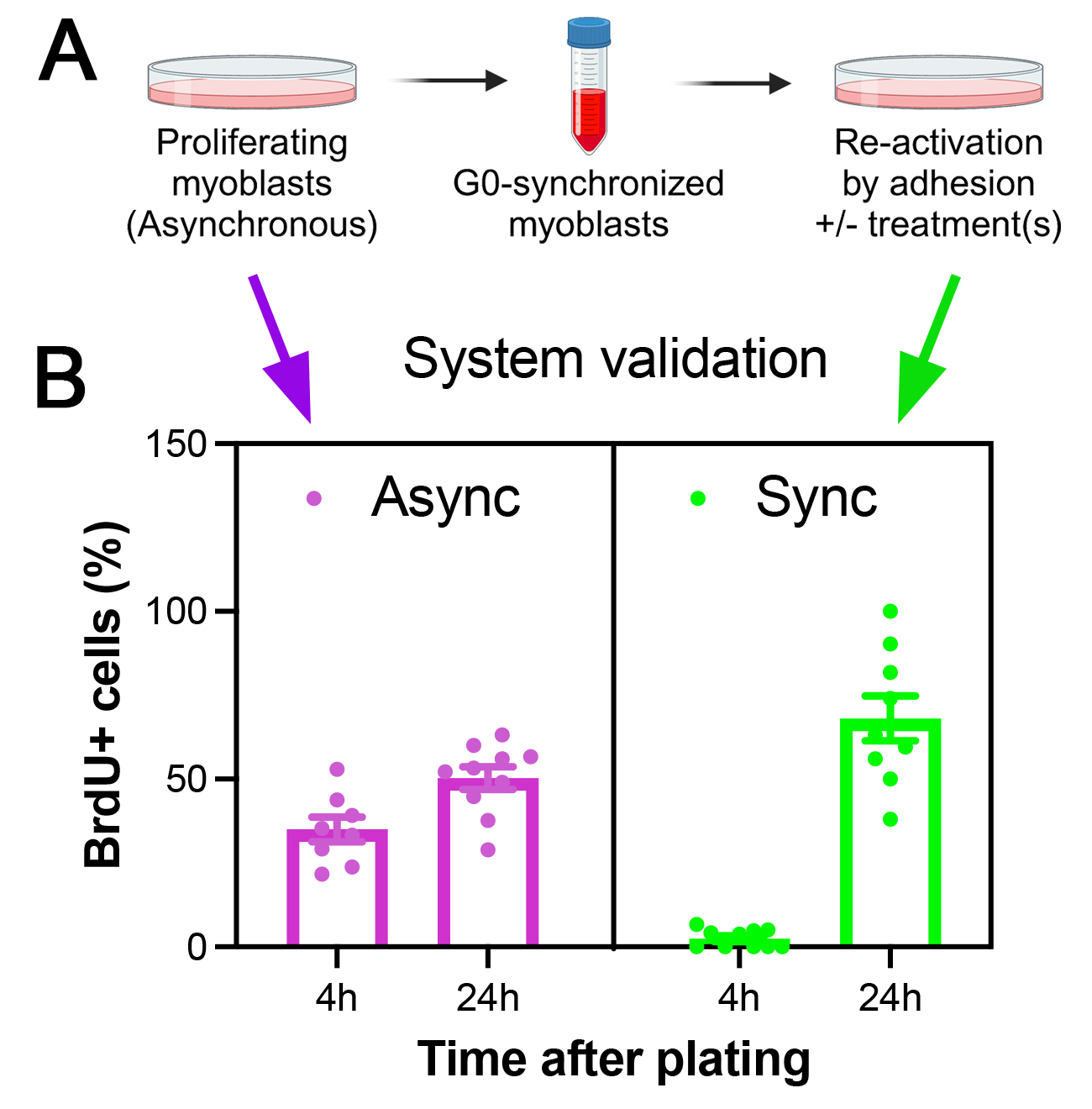
**Figure S1. Validation of the myoblast G0-synchronization method. A)** Diagram of the method. B) BrdU incorporation in asynchronous myoblasts 4 and 24 hours after plating (purple) and in G0-synchronized myoblasts also 4 and 24 hours after re-plating.

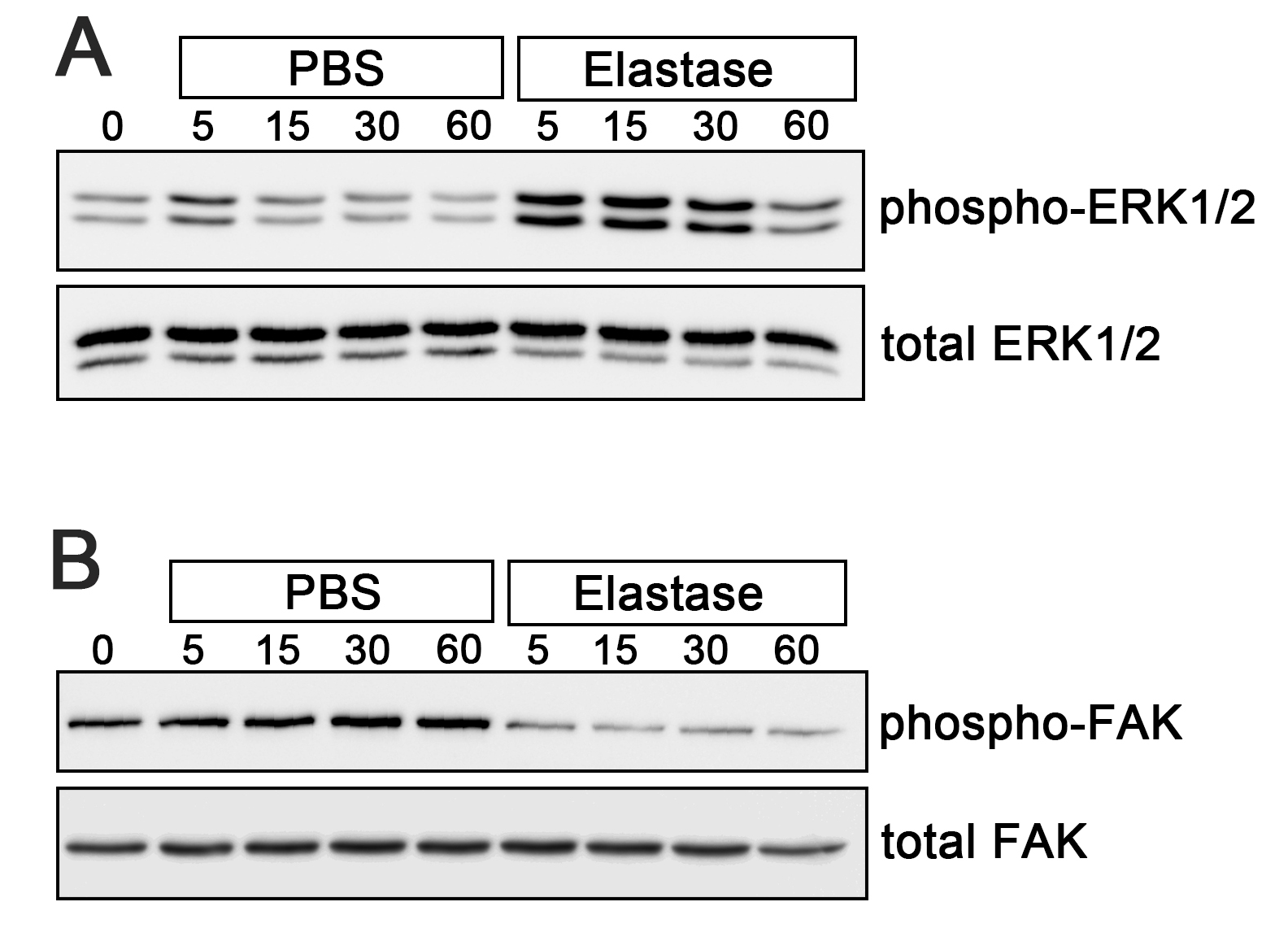

**Figure S2. Response to elastase stimulation is time-dependent. A)** ERK1/2 becomes phosphorylated in a time-dependent manner in response to elastase stimulation. **B)** FAK becomes immediately dephosphorylated in response to elastase stimulation and remains dephosphorylated for at least 1 hour.
